# ContiDesigner: Bioprocess Intensification through System-Level Design of Continuous Fermentation Cascades

**DOI:** 10.64898/2026.08.08.743657

**Authors:** Andrea C. Graf, Jürgen Zanghellini

## Abstract

Multi-stage continuous bioprocessing can increase volumetric productivity, operational consistency, and process throughput, but its design is complicated by coupling among dilution rate, reactor volume, feed allocation, and cellular physiology.

Here, we present ContiDesigner, available at https://chemnettools.anc.univie.ac.at/ContiDesigner/, a mechanistic steady-state framework and interactive web tool for the system-level design of continuous fermentation cascades. Comparing one- and two-stage configurations at equal total reactor volume and outlet flow, ContiDesigner reveals how internal flow and reactor volume allocation shape space-time yield and identifies productivity-maximizing operating conditions.

Compared with one-stage processes, two-stage cascades favor lower overall dilution rates, thereby preserving residence time in the production stage. The first-stage dilution rate approaches the corresponding one-stage productivity optimum, but the cascade optimum occurs earlier, reflecting a system-level compromise between biomass generation and production-stage residence time. However, two-stage operation outperforms optimized one-stage operation only when non-growth-associated production in the second stage is sufficiently strong, whereas increasing growth coupling favors one-stage operation.

Two case studies demonstrate both the potential and limits of process intensification. An optimized two-stage design is predicted to achieve a more than 1.5 fold increase in space-time yield for poly-R-3-hydroxybutyrate (PHB) production compared with a published experimental five-stage cascade, whereas the lactic acid case study identifies conditions under which staging offers no advantage. ContiDesigner translates these design principles into an accessible workflow to explore feasible operating regions and prioritize cascade designs for experimental evaluation.

**Graphical Abstract:** 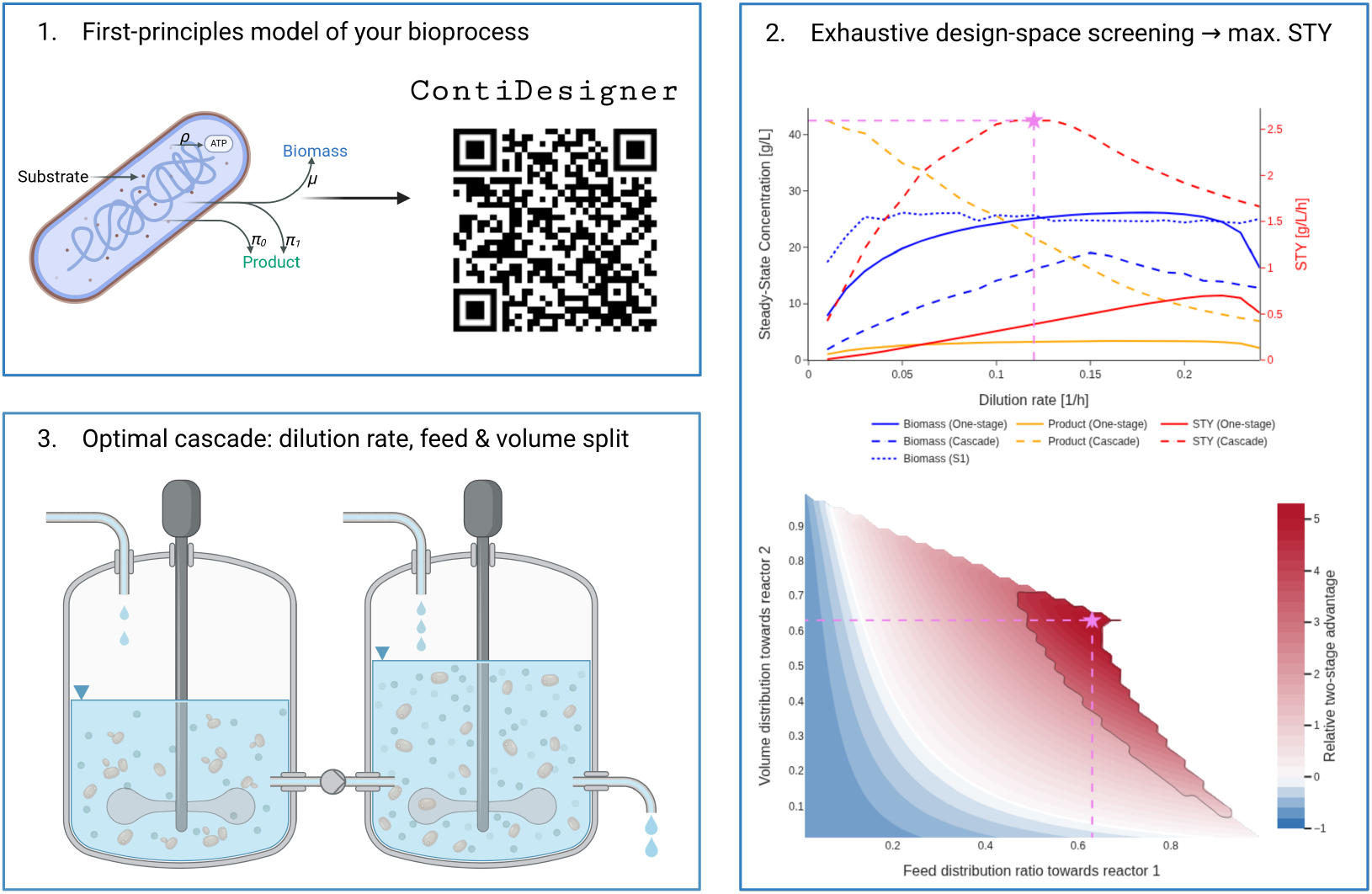

- `ContiDesigner` enables system-level design of continuous fermentation cascades
- High stage-one dilution supports biomass generation
- Low stage-two dilution preserves productive residence time
- Yet two-stage cascades favor lower overall dilution than one-stage systems
- Two-stage advantage requires strong non-growth-associated production in stage two

## 1. Introduction

The transition from petrochemical synthesis to microbial production is vital for achieving sustainable manufacturing goals [1]. Yet many biotechnological processes remain economically uncompetitive with chemical synthesis because of limitations in bioreactor performance, scale-up and process intensification [2, 3, 4]. Industrial bioprocesses still rely on batch and fedbatch operation, which suffer from variability, accumulation of metabolites, and downtime between runs [5, 6, 4]. Continuous production offers higher productivity, consistent quality and better facility utilization, but has seen limited large-scale adoption due to scale-up and control challenges, and the absence of accessible design frameworks [7, 5, 4].

One possibility to overcome these challenges is the use of multi-stage systems that separate growth and production phases, thereby mitigating inherent growth-production trade-offs and toxicity constraints [8, 9]. Classic studies on ethanol production showed that one-stage continuous fermentation could not exceed low titers because accumulating ethanol inhibited growth. In a two-stage design this limitation was alleviated by confining product accumulation to a downstream production reactor, allowing the upstream growth stage to remain protected from toxicity [10]. A more recent two-stage process for 1,3-propanediol production show that raising the dilution rate in stage 1 maximizes productivity in stage 1, but requires larger stage 2 volume to hit target titers [11]. These observations motivate cascaded designs and raise the question of how to distribute dilution and feed across stages to systematically optimize performance. Recent work has examined dilution rate optimization and feedback control in two-stage 1,3-propanediol fermentation, but broader design principles for allocating feed and dilution across stages remain limited [12].

Published work focuses mainly on optimizing cell physiology and control strategies [4, 13]. In contrast, reactor-level design still relies heavily on resource intensive empirical Design of Experiments (DoE) and, increasingly, black-box simulations and digital twin frameworks, which can accelerate optimization but often require substantial computational expertise and investment [14]. A shift towards knowledge-centric tools that explicitly encode mechanistic understanding and improve accessibility has been widely advocated [15, 16].

Mechanistic, first-principles models of continuous fermentation with various organisms are well established, including formulations that account for inhibition [17, 18, 19, 20] and multi-stage configurations [21, 22, 23, 11, 24]. While these models are valuable for describing specific systems, they rarely derive general design principles or explicitly separate growth generation from production specialization. Multi-stage studies often target niche objectives, such as substrate removal in wastewater treatment [25] or remain disconnected from practical, user-friendly design tools for production processes. At the same time, computational process modeling has been shown to substantially reduce experimental effort and overall development costs [15]. To our knowledge, no open, general-purpose framework focusing specifically on the design of cascaded two-stage continuous production processes is available, and we are not aware of any that combine steady-state analysis with design-space exploration for such systems.

This work addresses these gaps with a mechanistic steady-state analysis of one- and two-stage continuous bioprocesses under Monod growth [26] and Luedeking-Piret production kinetics [27]. We implement this as an open-access web tool that visualizes feasible operating regions, trade-offs, and near-optimal designs, allowing users without computational expertise to compare one-vs. two-stage setups. It prioritizes process understanding over black-box optimization by explicitly revealing the underlying mechanistic relationships and helps focus experimental efforts on the most promising regions of the design space.

## 2. Methods

### 2.1. Model scope and process configurations

We consider continuous microbial production processes operated either as a single continuous stirred tank reactor (CSTR) (one-stage process) or as two CSTRs arranged in series (two-stage process; Fig. 1). All reactors operate at constant liquid volume, with perfect mixing and without cell retention. The model is intended as a system-level design framework for processes in which washout, biomass generation, and volumetric productivity are the dominant design constraints [5]. Model notation is summarized in Supplementary Information Tab. A.1.

**Figure 1:**
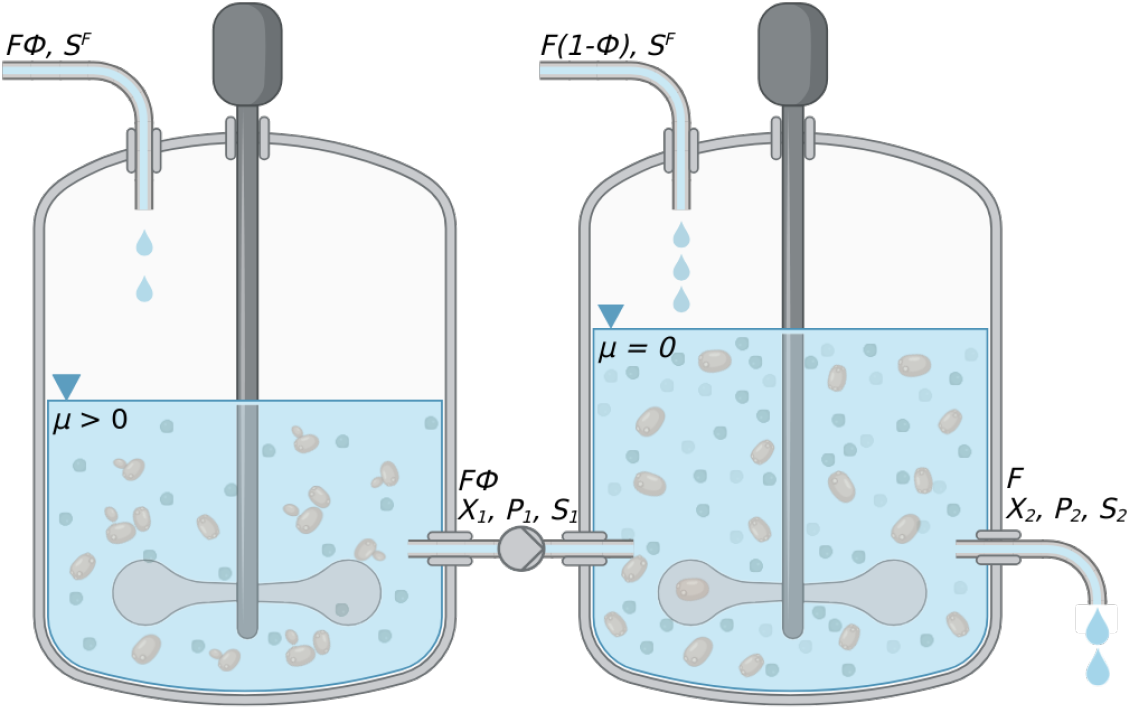
Schematic representation of a two-stage chemostat cascade where the first reactor receives only fresh medium and supports cell growth. The second reactor operates in a growth arrested regime and is supplied with the outflow from the first stage as well as an additional feed of fresh medium.

To compare a one-stage process and a two-stage process on an equal basis, both configurations are assigned the same total reactor volume *V* and the same final outlet flow rate *F* . For the two-stage process, these are given by *V* = *V*_1_ + *V*_2_, where *V*_*i*_ denotes the reactor volume of stage *i. F*_1_ denotes the outlet flow of stage 1, and the second, final outlet flow of the cascade is *F*_2_ = *F* . The additional fresh feed into stage 2 is therefore *F*_2_ −*F*_1_. With this notation, the one-stage process is recovered as the limiting case *V*_2_ = 0 and *F*_2_ = *F*_1_ = *F* . The resulting overall dilution rate is therefore

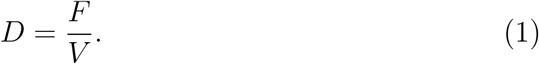

For the one-stage process, this gives *D*_OS_ = *F*_1_*/V*_1_ = *D*. For the two-stage process, *D* describes the overall process dilution rate rather than the local dilution rate *D*_*i*_ in either individual reactor *i*.

To express the local dilution rates of the two-stage in terms of *D*, we define the flow fraction *ϕ* and the volume fraction *ν*,

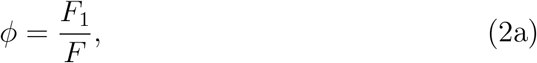

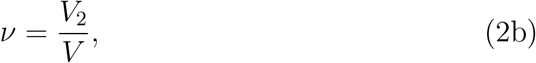

respectively. The corresponding local dilution rates are then

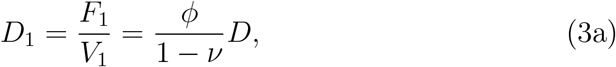

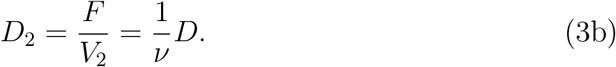

Here, *D*_1_ is the dilution rate of stage 1, whereas *D*_2_ is the dilution rate of stage 2, whose outlet flow is the final cascade flow *F* = *F*_2_. Thus, *D* controls the overall process intensity, while *ϕ* and *ν* determine the internal distribution of flow and reactor volume. The one-stage process is treated as a separate model and corresponds formally to the limiting case *ϕ* →1 and *ν* →0. However, quantities associated with stage 2 are not defined in this limit.

This comparison fixes total reactor size and overall process throughput. Differences in predicted performance therefore arise from the internal distribution of volume and flow, not from differences in total reactor volume or final outlet flow.

Unless stated otherwise, both fresh-feed streams are assigned the same substrate concentration, 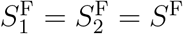. The total substrate feed rate in the two-stage process is therefore

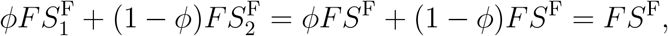

matching the one-stage process. However, in process-intensification scenarios, the stage 2 feed concentration is allowed to exceed the baseline feed concentration subject to 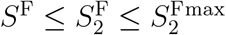.

### 2.1.1 Reactor model

Each reactor *i* is described by mass balances for biomass *X*_*i*_, product *P*_*i*_, and limiting substrate *S*_*i*_ (Fig. 1):

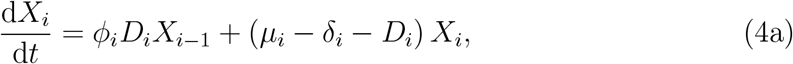

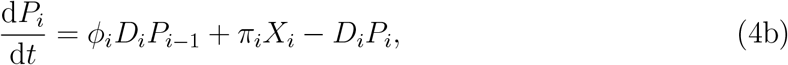

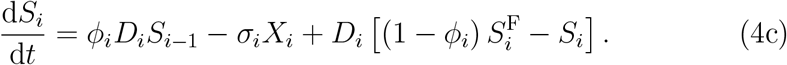

Here, *µ*_*i*_ and *δ*_*i*_ are the specific growth and death rates, respectively, *D*_*i*_ = *F*_*i*_*/V*_*i*_ is the local dilution rate, and *ϕ*_*i*_ = *F*_*i*−1_*/F*_*i*_ is the fraction of the inlet flow to reactor *i* originating from the preceding reactor. The remaining fraction 1 −*ϕ*_*i*_ enters as fresh feed with substrate concentration 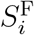. Throughout the analysis, we take the death rate to be stage-independent, *δ*_*i*_ = *δ* for all *i*.

For the first reactor, there is no upstream reactor, so *F*_0_ = 0 and *ϕ*_1_ = 0.

The one-stage process mass balance is recovered for *i* = 1, *ϕ*_1_ = 0, and *D*_1_ = *D*. In the two-stage process, stage 1 is described by *i* = 1, *ϕ*_1_ = 0, and 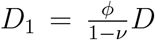. The stage 2 balance is obtained for *i* = 2, *ϕ*_2_ = *ϕ*, and 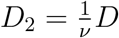, as defined in (3).

### 2.1.2. Cellular model

The specific growth rate in reactor *i* is described by

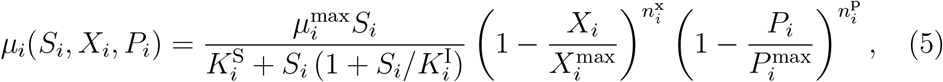

where 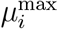 is the maximum specific growth rate, 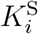 is the Monod half-saturation constant, 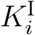 is the substrate inhibition constant, and 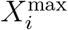 and 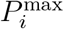are the biomass and product inhibition thresholds, respectively, and 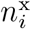 and 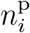 are empirical inhibition exponents controlling the strength of biomass and product inhibition [28, 29]. Note that the inhibition thresholds 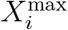 and 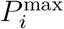need to be chosen such that 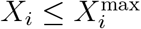 and 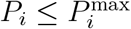 over the simulated concentration range, ensuring non-negative inhibition factors. Standard Monod kinetics are recovered for 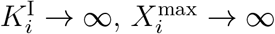, and 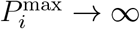 Product formation follows Luedeking–Piret kinetics,

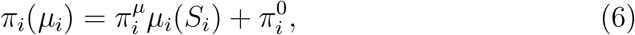

where 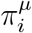 and 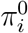 are the growth-associated production coefficient and non-growth-associated production rate in reactor *i*, respectively.

Substrate consumption combines contributions from growth, product formation, and maintenance:

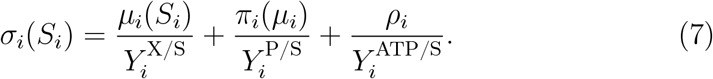

Here, 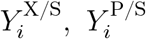, and 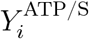 denote the maximum theoretical biomass, product, and ATP yields on substrate, respectively, and *ρ*_*i*_ is the specific maintenance ATP requirement. This formulation treats substrate demand for growth, product formation, and maintenance as additive, without explicitly modeling intracellular allocation or regulatory trade-offs between these processes.

In the following, we assume that stage 1 supports growth according to (5), whereas growth in stage 2 is arrested, *µ*_2_ = 0. However, the software implementation also allows incomplete growth arrest with 0 ≤ *µ*_2_ *< µ*_1_.

### 2.2. Design space exploration

Process performance is evaluated by the space-time yield,

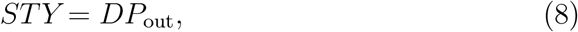

where *P*_out_ is the product concentration in the final outlet stream. For the one-stage process, *P*_out_ = *P*_OS_ , whereas for the two-stage process, *P*_out_ = *P*_2_.

For the one-stage process, *STY* is optimized over the overall dilution rate *D*:

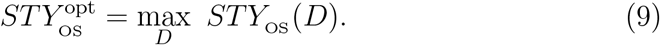

For the two-stage process, the optimization additionally includes the flow and volume allocation variables *ϕ* and *ν*:

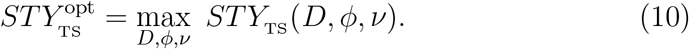

All optimizations are restricted to feasible operating points. For the one-stage process, feasibility requires non-washout operation,

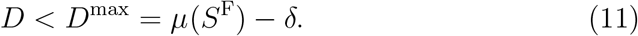

For the two-stage process, the same condition applies to stage 1 with its local dilution rate,

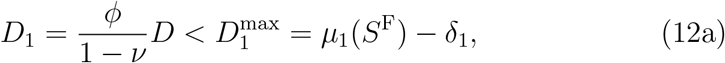

with admissible allocation variables 0 *< ϕ*≤ 1 and 0 *< ν <* 1. Because biomass is supplied by stage 1, stage 2 does not impose an additional upper washout bound on *D*. Instead, solving the stage 2 biomass balance at steady state gives

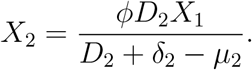

Thus, a positive finite stage 2 biomass concentration requires *D*_2_+*δ*_2_ − *µ*_2_ *>* 0, or equivalently

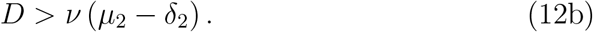

For the growth-arrested base case, *µ*_2_ = 0, this condition is automatically satisfied for any *D >* 0.

The resulting feasible design space is then evaluated for both configurations to identify the maximum attainable *STY* under the specified feed scenario.

### 2.3. Implementation

For each specified set of process constraints and physiological parameters, the dynamic mass-balance equations (4), together with the kinetic definitions in (5)–(7), are evaluated at steady state by setting the time derivatives to zero and solving the resulting algebraic equations for the one-stage process and two-stage process. The resulting steady-state concentrations, provided in the Supplementary Information (Eqs. (A.1) to (A.3)), are then used to compute *STY* according to (8) over the feasible design space defined by the dilution-rate relations (1) and (3), the allocation constraints 0 *< ϕ* ≤ 1 and 0 *< ν <* 1, and the feasibility conditions (11), and (12).

The model is implemented in Python as an object-oriented class structure. Source code and a Dash/Plotly ContiDesigner webtool (hosted at https://chemnettools.anc.univie.ac.at/ContiDesigner/) are available at https://github.com/acgraf/ContiDesigner under the MIT license. Users specify process constraints, including total reactor volume and feed substrate concentrations, as well as physiological parameters, including Monod kinetics, maximum theoretical yields, and Luedeking–Piret coefficients, which can be estimated from literature data or shake-flask experiments.

The workflow comprises:

1. At each dilution rate *D*, evaluating the steady-state performance of the one-stage process and identifying the optimal two-stage process configuration over (*ϕ, ν*).
2. Computing the two-stage process *STY* landscape over (*ϕ, ν*) at selected values of *D*.
3. Simulating the time evolution of the one-stage process and two-stage process configurations.
4. Exporting steady-state results, optimal designs, and model definitions as `CSV` and `SBML` files. The exported `SBML` files are compatible with `SBML`-based modeling tools, including `COPASI` [30], `sbmlutils` [31], and `Tellurium` [32].

The web tool enables interactive exploration of these results. Users select operating points by clicking dilution-rate curves or contour points. Changing *D* triggers recomputation of the (*ϕ, ν*) grid, and the interface highlights both the global optimum and the selected operating point. Users can also export the corresponding results as tables. An overview of the web interface is shown in Fig. 2.

**Figure 2:**
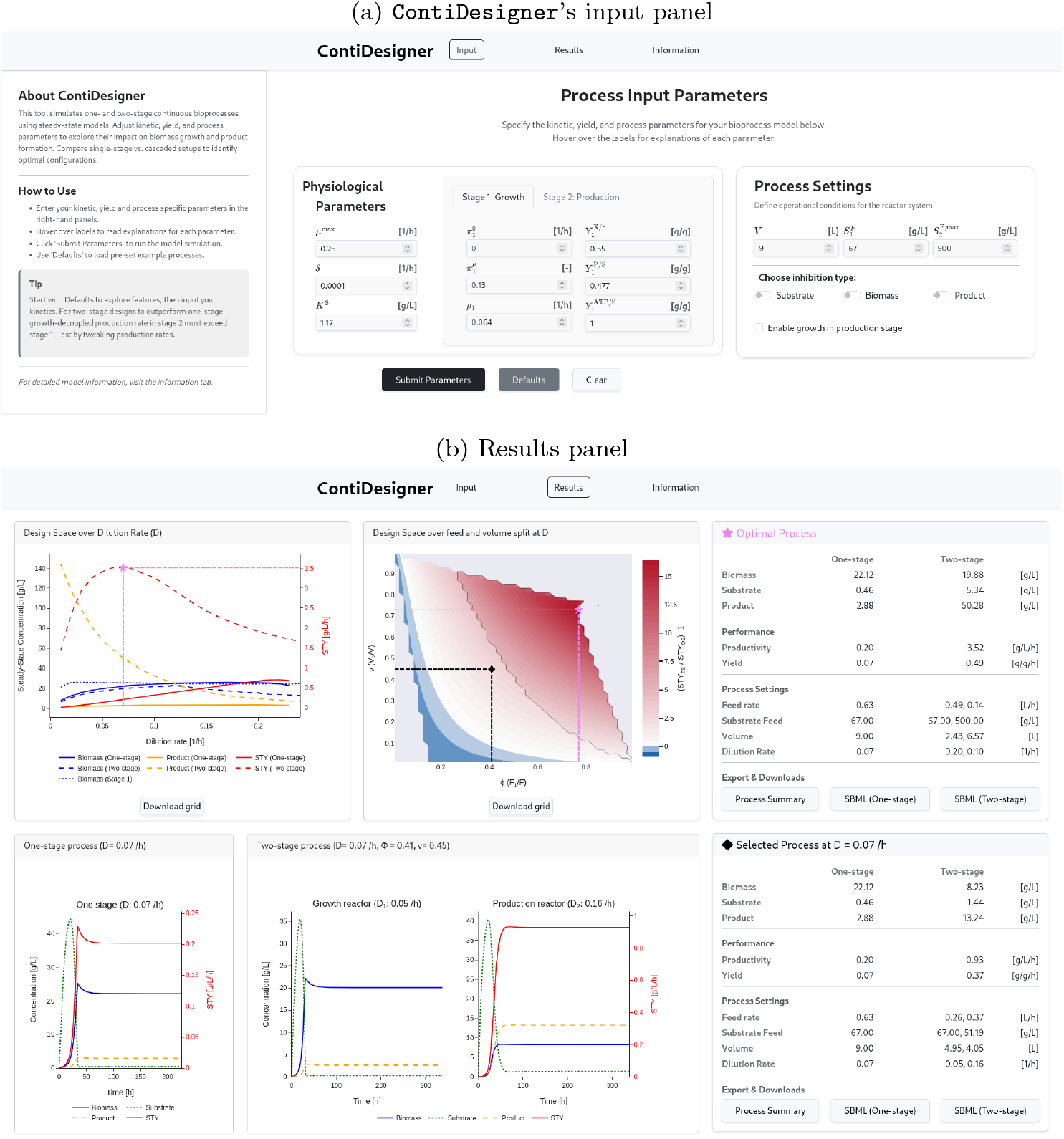
Interface of ContiDesigner. (a) Input panel for kinetic parameters, process settings, and stage-specific options, including inhibition and arrested growth. The parameters shown correspond to the PHB case study. (b) Interactive results panel. The upper panels display steady state behavior across dilution rates and the feed and volume ratios (*ϕ* and *ν*) at the selected dilution rate. The pink star marks the global optimum and the black diamond the selected operating point. The contour plot additionally highlights the region of process intensification, indicated by increased color saturation, where the second-stage feed concentration exceeds that of the first stage 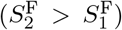. The lower panels show dynamic simulations of the one- and two-stage processes and summary tables for the optimal and selected designs

### 2.4. Case studies

Case-specific model parameters and operating conditions are provided in the Supplementary Information (Tab. A.2) and included as default configurations in ContiDesigner.

### 2.4.1. PHB

We implemented the five-stage CSTR cascade for PHB production by *Cupriavidus necator* described by Atlić et al. [33], using Horvat et al.’s [21] kinetic parameters. Reactor volumes, flow rates, and substrate feed concentrations were taken from the original experimental setup.

The experimental process relies on nitrogen limitation to induce PHB accumulation. We do not model nitrogen limitation explicitly, but approximate phenomenologically by assuming full growth kinetics in R1, growth-limited transition in R2, where *µ*_2_ ≈ 0.0125 h^−1^ (estimated from steady state biomass) and growth arrest in R3-R5. Substrate feeds were fixed at reported values. Unlike Horvat et al., who model the non-growth-associated PHB production rate as substrate dependent, our model assumes production rate to be constant. Therefore, PHB production was set to zero once the substrate concentration approached depletion (*s* → 0). Both *µ*_2_ = 0.0125 h^−1^ and *µ*_2_ = 0 h^−1^ formulations were tested.

For the alternative two-stage process, growth was arrested in the second stage, while the total reactor volume and cellular kinetic parameters were kept identical to those of the five-stage cascade. The overall dilution rate *D*, flow allocation *ϕ*, and volume allocation *ν* were optimized. The optimization was repeated at different fixed total substrate feed rates.

### 2.4.2. Lactic acid; necessary production improvement

We adapted the steady state one-stage lactic acid fermentation model reported by Gordeeva et al. [17]. The original model describes biomass, substrate, lactic acid, and a precursor pool *M* , and investigates the effects of substrate, biomass, and product inhibition on steady states over a range of dilution rates.

Two modifications were required to represent the published model within our framework.

1. In the original model, substrate is consumed for biomass growth but not for lactic acid production. For benchmark reproduction, we therefore set the product yield to *Y* ^P/S^ = 100 g g^−1^, making the additional substrate demand associated with product formation negligible.
2. Gordeeva et al. [17] define an original substrate-feed concentration of *s*^F^ = 60 g L^−1^ and a precursor concentration of *M*_0_ = 20 g L^−1^. The precursor is converted to substrate with a rate constant of *k*_*M*_ = 0.035 h^−1^. In our implementation, the precursor pool was represented by adding its steady-state contribution to the original substrate-feed concentration:

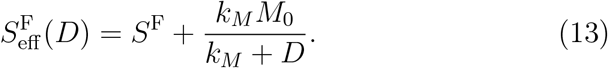

With these adaptations and a fixed effective substrate-feed concentration of 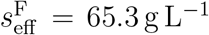 , evaluated at *D* = 0.1 h^−1^, our implementation closely reproduced the steady-state profiles reported by Gordeeva et al. (Fig. A.6).

For the subsequent comparison of the one-stage process and two-stage process, we omitted the precursor pool by setting *M*_0_ = 0, fixed *S*^F^ = 60 g L^−1^, and replaced the benchmark yield with the theoretical maximum product yield, *Y* ^P/S^ = 1 g g^−1^, for the conversion of glucose to lactic acid. The optimized one-stage process and two-stage process were compared using the relative difference in optimal *STY*,

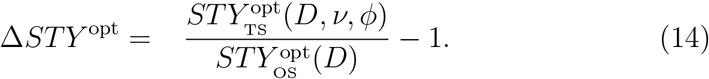

## 3. Results

### 3.1. Case studies

#### PHB

To evaluate our reactor model, we first compared it to the five-stage PHB cascade reported by Atlić et al. [33]. In this nitrogen-limited process, biomass formation occurs mainly in the early reactors, whereas PHB accumulates across the downstream production reactors. Using the mass-balance formulation in (4) together with the kinetic definitions in (5)–(7), our implementation captured the major biomass and PHB accumulation trends (Supplementary Information, Tab. A.3). The corresponding dynamic concentration profiles across the five-reactor cascade are shown in Fig. A.5.

We next asked whether a simplified two-stage process could recover the performance of the published five-stage cascade. We therefore used ContiDesigner to design a two-stage configuration with a growth-supporting first stage and a growth-arrested production stage. The ContiDesigner interface enables definition of process parameters and kinetic constraints, followed by interactive exploration of optimized operating conditions (Fig. 2).

The optimized two-stage design (Fig. 3) operated at *D* ≈ 0.07 h^−1^, more than twice the overall dilution rate of the original five-stage cascade (see Supplementary Information, Tab. A.3). It used *D*_1_ = 0.2 h^−1^ and *D*_2_ = 0.1 h^−1^, with *F*_1_ = 0.49 L h^−1^, *F*_2_ = 0.63 L h^−1^, *V*_1_ = 2.43 L, and *V*_2_ = 6.57 L. Despite reaching a lower PHB titer than the reported experimental cascade (53 g L^−1^ vs. 63 g L^−1^), it predicted to improve *STY* by more than 1.5 fold, reaching 3.52 g L^−1^ h^−1^.

**Figure 3:**
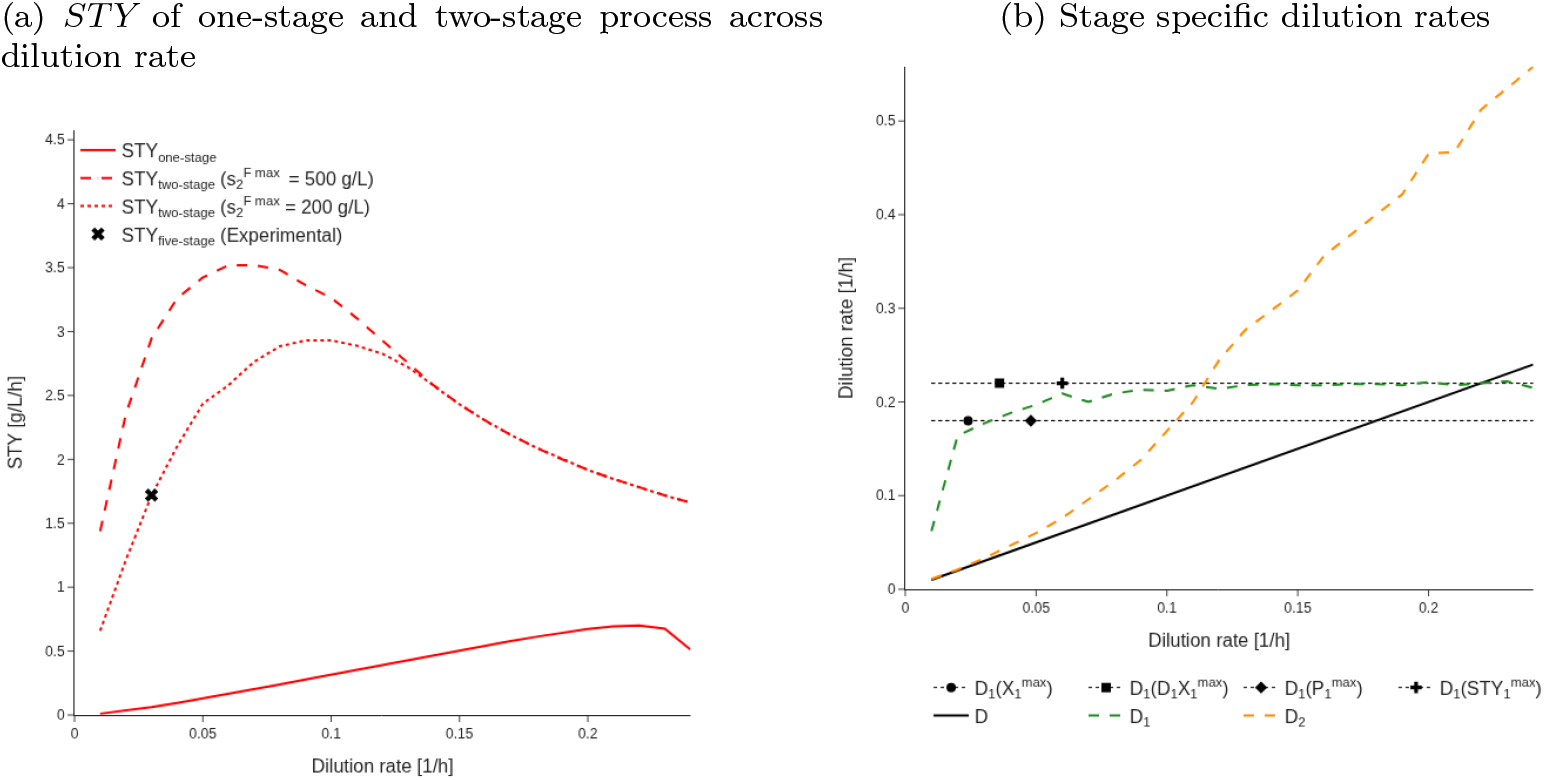
Performance of the one-stage optimum and the optimized two-stage cascade across the overall dilution rate *D*, using kinetic parameters from the PHB case study [21]. Panel (a) shows one-stage and two-stage *STY*. Experimental results reported by Atlic et al. are indicated by a black “x”. The experimental five-reactor cascade operated at an overall dilution rate of *D* = 0.03 h^−1^. This configuration can be reduced to a two-stage cascade while achieving higher *STY*, with an optimum at *D* = 0.07 h^−1^. A two-stage process constrained to achieve the same *STY* as the experimental setup (dotted red line) at *D* = 0.03 h^−1^ would require a maximum substrate concentration in stage 2 of 200 g L^−1^. This is lower than the 500 g L^−1^ substrate concentration used in the experimental feed of reactors R2-R5 and reflects the substrate limitation imposed by the experimental feeding strategy. Panel (b) shows the optimized dilution rates *D*_1_ and *D*_2_ as functions of *D. D*_1_ saturates at one-stage optimum *D* ∼ 0.22 h^−1^ and does not exceed it. Only when *D* itself is larger than the one-stage optimum, i.e. as the system approaches washout, does *D* become larger than *D*_1_. *D*_2_ increases with *D* but more rapidly, remaining above *D* throughout. The horizontal lines indicate the dilution rates in stage 1 when maximizing *X*_1_ and *P*_1_, and those maximizing throughput rates *STY*_1_ and *X*_1_ · *D*_1_. For this case study, where production in stage 1 is fully growth coupled, the optima of *X*_1_ and *P*_1_ coincide, as do the optima of *STY*_1_ and *X*_1_*D*_1_.

To better understand the optimized two-stage process behavior, we compared it with an equivalent one-stage process using the same total volume, outlet flow, and kinetic parameters (Fig. 3). This comparison provides a single-reactor reference for interpreting the optimized local dilution rates (*D*_1_ and *D*_2_) relative to the overall dilution rate *D*. The optimized two-stage process rapidly drives *D*_1_ according to the competing requirements of biomass formation and production-stage operation. The resulting values of *D*_1_ remain within the range defined by the intrinsic optima of stage 1, at the dilution rates maximizing *X*_1_, *P*_1_, *STY*_1_ and biomass throughput rate (*X*_1_*D*_1_). For the present case study, in which production in stage 1 is fully growth-coupled, these optima coincide pairwise, such that biomass and product concentration are maximized at one dilution rate, while *STY*_1_ and *X*_1_*D*_1_ are maximized at a higher dilution rate. However, this relationship is not generally expected when the distribution between growth-coupled and growth-decoupled production is varied.

At low overall dilution rates, the optimized cascade operates with relatively low *D*_1_, reflecting the limited substrate supply to stage 2 and the need to maintain sufficient residence time in the production stage. At low *D*, this adjustment is limited by the total substrate supply. As *D* increases, dilution in stage 1 increases and approaches the dilution rate maximizing first-stage productivity. However, the optimum cascade performance is reached before this upper limit is attained: at the optimal overall dilution rate, *D*_1_ is still increasing toward the stage 1 productivity optimum. Throughout the investigated range, *D*_2_ *> D*, as required by *D*_2_ = *D/ν* with 0 *< ν <* 1, see Eq. (3b). Initially, *D*_2_ remains only slightly above the overall dilution rate. Once *D*_1_ approaches the upper stage 1 productivity limit, *D*_2_ begins to diverge more strongly from *D*, shortening the production-stage residence time *τ*_2_ = 1*/D*_2_ and thereby decreasing *STY*.

From the analysis of this case study, a design rule emerges: the two-stage process favors the lowest overall dilution rate that still allows *D*_1_ to approach the equivalent one-stage process optimum, thereby preserving residence time in stage 2 without compromising biomass export from stage 1. This is the opposite of the one-stage process, for which optimal operation requires increasing the dilution rate toward its productivity maximum.

#### Lactic acid

The PHB case demonstrates that stage separation can substantially improve productivity. However, is a two-stage process always superior to an optimized one-stage process? To answer this question, we use lactic acid production as a model system and systematically vary how production capacity is partitioned between growth-coupled and non-growth-associated production and between the two stages. We characterize these allocations using two state-independent reference measures. Specifically, we define the fraction of the production capacity in stage 1 that is growth-coupled at the reference growth rate *µ* = *µ*^max^,

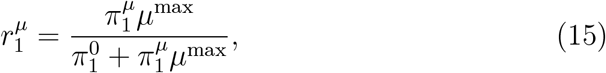

and the ratio of non-growth-associated production capacity in stage 1 to that in stage 2,

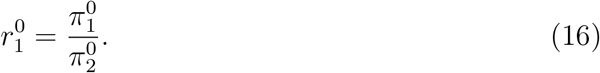

Here, 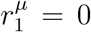 and 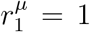 denote purely non-growth-associated and purely growth-coupled production in stage 1, respectively. Similarly, 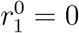 assigns all non-growth-associated production capacity to stage 2, whereas 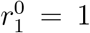 distributes capacity equally to stage 1 and stage 2. Cases with 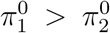 are not considered since this would correspond to a decrease in production capacity upon transition to stage 2.

Unlike the state-independent reference measure 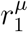, the realized growth-coupled production fraction depends on the steady-state growth rate 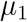 and can only be determined after solving the reactor model. Evaluating the ratio at *µ*^max^ therefore provides a fixed parameterization that can be varied independently of the resulting process state.

Using these measures, Fig. 4a maps the relative change in optimal *STY* of the two-stage process compared with the one-stage process over 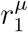 and 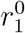. Positive values indicate that the two-stage process outperforms the one-stage process, whereas values close to zero indicate that staging provides no benefit and the optimal two-stage process effectively reduces to the one-stage process.

**Figure 4:**
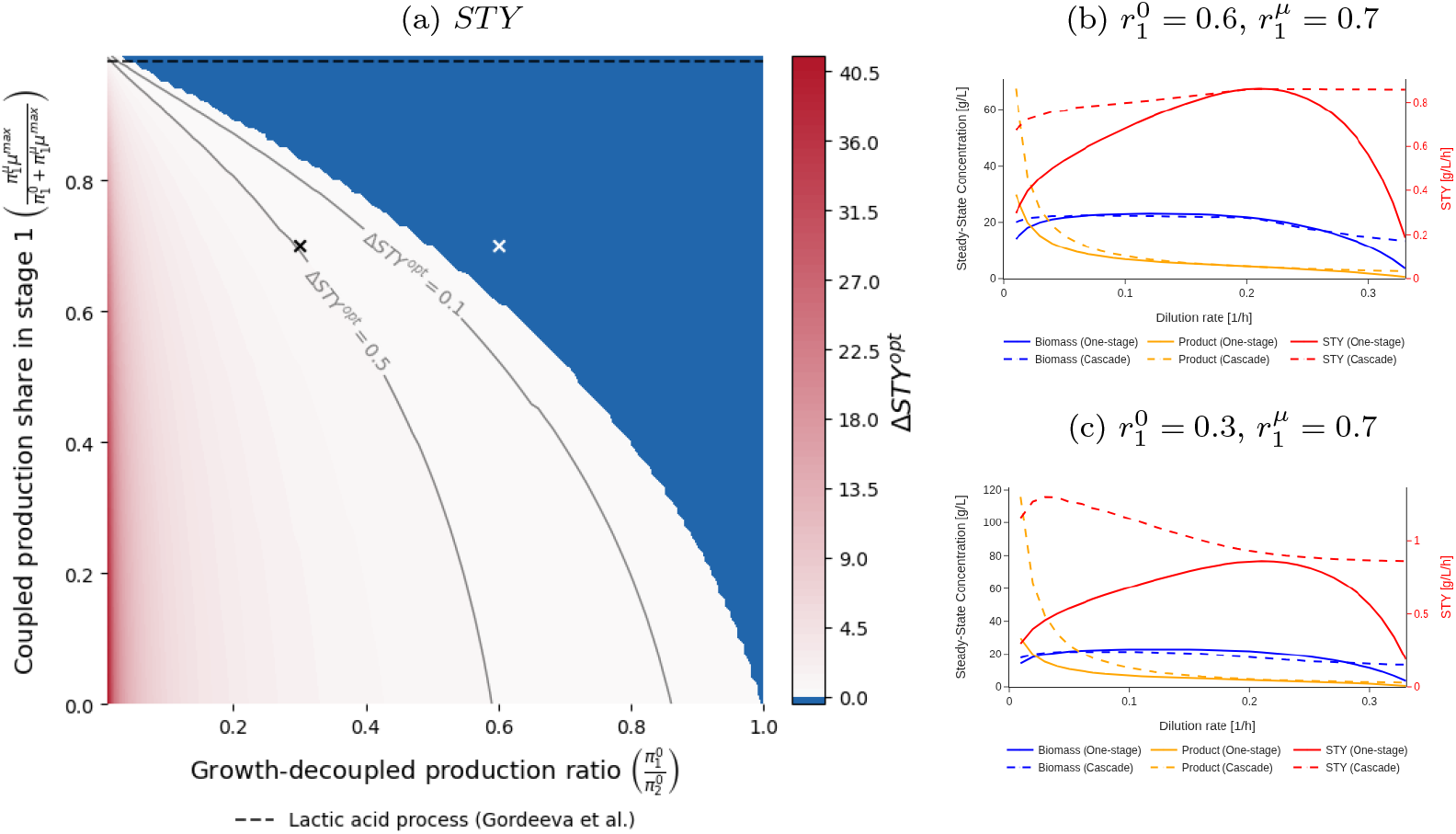
Lactic acid: (a) Optimal productivity advantage of the two-stage process relative to the optimal one-stage configuration, Δ*STY* ^opt^, across state-independent production allocation fractions 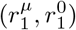. The horizontal dashed line corresponds to the reference lactic acid process of Gordeeva et al. This process requires 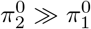, only at 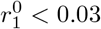 is the two-stage advantageous, yielding only∼ 10% *STY* improvement even at 38.5× higher 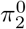. High 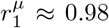 biases toward one-stage. Panels (b) and (c) show the corresponding dilution ranges and optimal steady states for two-stage and one-stage where (b) illustrates the region favoring one-stage and (c) the region favoring two-stage. In regions where two-stage outperforms one-stage (c), the one-stage optimum lies at higher dilution 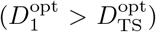. In one-stage-favoring regions (panel b), the one-stage optimum also exceeds the two-stage optimum 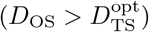.

For the lactic acid case study based on Gordeeva et al. [17], Fig. 4a shows that an optimized two-stage process rarely outperforms the one-stage process. This is because lactate production in the reference process is highly growth-coupled, with 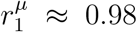 (horizontal dashed line in Fig. 4a). At this value, the two-stage process becomes advantageous only for 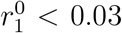 Increasing the production rate in stage 2 to 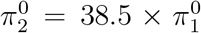 results in an improvement of only about 10 % compared with the one-stage process. Thus, strong growth coupling favors the one-stage process, whereas a two-stage process becomes beneficial only when the non-growth-associated production capacity in stage 2 is substantially greater than that in stage 1.

## 4. Discussion

Continuous bioprocessing offers the potential for high volumetric productivity, consistent product quality, and efficient reactor utilization [34, 35]. Realizing these benefits, however, requires process configurations in which growth, production, and residence time are coordinated at the system level [36, 37].

An intuitive design goal is to combine high biomass generation in stage 1 with a long residence time for production in stage 2. Our results, however, show that these operating states cannot be fully decoupled (Fig. 3b). Because the stages remain connected through the transfer of biomass, substrate, and product, the optimized cascade exhibits only a partial division of labor. The overall optimum is therefore a coordinated compromise between biomass generation in stage 1 and productive residence time in stage 2, rather than a combination of the individual stage optima.

The parameter-space analysis in Fig. 4a revealed a clear threshold separating regimes in which the one-stage process or two-stage process is optimal. The two-stage process outperforms the one-stage process only when the non-growth-associated production capacity in stage 2 is sufficient to compensate for both the growth-associated production available in the one-stage process and the coupled allocation of reactor volume, flow, and residence time between the two stages. Consequently, the required production advantage of stage 2 increases as growth coupling in stage 1 becomes stronger.

For a two-stage process, strain design should therefore explicitly target high non-growth-associated production in stage 2, rather than optimize primarily for growth-coupled production. SimulKnock [37], for example, maximizes biomass formation at the cellular level and may therefore favor growth-supporting phenotypes suited to one-stage process operation rather than explicitly exploring a separate growth-arrested production state. High non-growth-associated production alone may nevertheless be insufficient if substrate uptake declines after growth arrest, as shown by Klamt et al. [8]. They proposed enforced ATP wasting to increase non-growth-associated maintenance demand and thereby sustain substrate uptake during production. When ATP generation is coupled to product formation, this strategy can support production in the non-growing state. Such a mechanism can be represented in our framework by linking 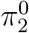 to the cellular maintenance requirement.

The published PHB cascade [33, 21] provides a practical benchmark for the system-level trade-offs identified here. With the stage 2 feed concentration limited to the experimentally used value of 500 g L^−1^, the optimized two-stage configuration was predicted to achieve a 1.6*×* higher *STY* than the experimentally implemented five-stage cascade. This improvement arose from reallocating reactor volume and increasing the overall dilution rate while maintaining sufficient residence time in the production stage. Conversely, at the overall dilution rate of the experimental cascade, an optimized two-stage configuration was also predicted to reproduce the reported *STY* with a maximum stage 2 feed concentration of approximately 200 g L^−1^. This indicates that optimized flow and volume allocation can reduce the feed concentration required to meet a specified productivity target.

These findings are relevant because productivity and process design remain important challenges in microbial PHB production [38]. Alternative feedstocks may improve raw-material costs and environmental performance, but PHB production remains economically challenging relative to petrochemical plastics [39, 40]. Although the present framework does not evaluate process economics directly, it addresses one contributor to economic performance by identifying how reactor architecture and operating conditions constrain attainable volumetric productivity.

Although the present framework focuses on two-stage operation, the same system-level coupling applies to cascades with additional reactors. Additional stages may facilitate physiological transitions, improve process control, or provide greater operational robustness. However, our results show that an optimized two-stage configuration can achieve a *STY* comparable to that of the experimentally implemented five-stage PHB cascade. This is consistent with previous optimization results showing that the benefit of adding further reactors may diminish rapidly. For a fermentation model with simultaneous substrate and product inhibition, Abu Reesh identified two to three optimally sized reactors as the best configuration for minimizing total reactor volume [41]. Nevertheless, the optimal number of stages depends on the assumed kinetics and optimization objective, and a direct comparison with an independently optimized multi-stage PHB cascade remains open.

Existing optimization frameworks such as OptMSP [36] provide powerful programmable approaches for multi-stage bioprocess optimization, but require users to formulate and implement the underlying process model. In contrast, ContiDesigner is designed for rapid exploration of continuous fermentation cascades through an interactive workflow that integrates reactor partitioning, interstage flow allocation, and steady state design-space visualization.

ContiDesigner operationalizes these insights via FAIR [42] multi-level access: interactive design-space plots, `SBML`/`CSV` exports, with extensible Python core. Rather than black-box optimization, it exposes feasible regions and trade-offs. That makes the framework useful for both modelers and experimentalists who want to connect kinetic assumptions to concrete cascade configurations.

## 5. Conclusion

Steady state analysis of two-stage continuous bioprocesses show that system-level design is essential, because stage-wise optimization does not generally identify the global productivity optimum. The results yield design rules showing that stage 1 should operate near the biomass throughput maximum of the one-stage system while stage 2 should maximize the substrate feed concentration and minimize additional feed volume. Two-stage cascades outperform an optimized one-stage operation only when stage 2 dominates non-growth-associated production, and even then they must overcome the inherent engineering penalties of splitting the system. ContiDesigner translates these insights into an open, interactive web tool that enables rational process design from basic kinetic data, bridging mechanistic understanding and practical experimentation.

## Declarations

### Ethics approval and consent to participate

Not applicable.

### Declaration of AI use

During the preparation of this work, the authors used generative AI to assist with code development and debugging and to improve the manuscript’s language, readability, and flow. All AI-assisted text and code were reviewed, edited, and validated by the authors, who take full responsibility for the manuscript, analyses, and results.

### Consent for publication

We confirm that the manuscript has been read and approved by all named authors and that there are no other persons who satisfied the criteria for authorship but are not listed. We further confirm that the order of authors listed in the manuscript has been approved by all of us.

For open access purposes, the authors have applied a [CC BY public copyright license](https://creativecommons.org/licenses/by/4.0/) to any author-accepted manuscript version arising from this submission.

### Code and Data availability

All source code, model files, and data underlying this study are freely available at https://github.com/acgraf/ContiDesigner under the [MIT license](https://opensource.org/license/mit). An archived version with a persistent DOI will be deposited in Zenodo upon acceptance.

### Materials availability

Not applicable

### Author contribution

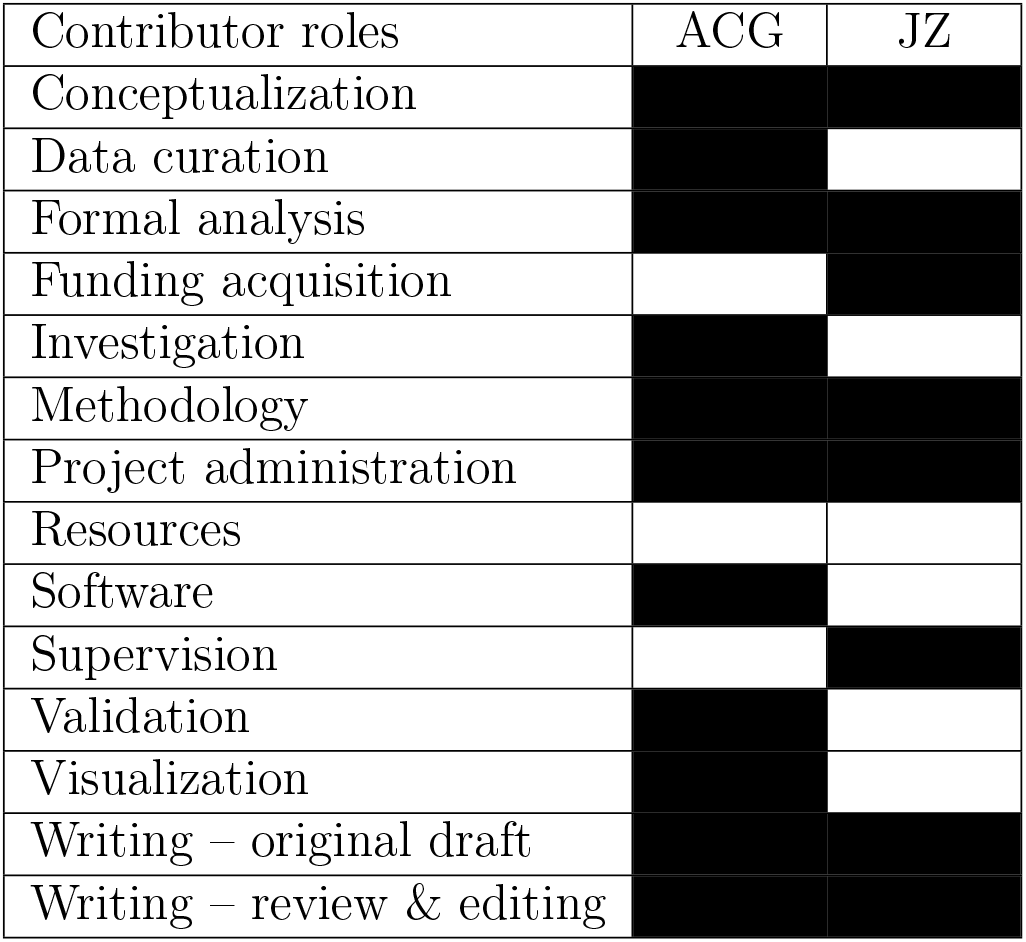

