## Supplementary Information for "ContiDesigner: Bioprocess Intensification through System-Level Design of Continuous Fermentation Cascades"

### Appendix A. Supplementary Information

#### *Appendix A.1. Model notation*

Table A.1: Model notation. Subscript $i$ denotes the reactor stage, with $i = 1, 2$ corresponding to stage 1 and stage 2, respectively. Model input parameters are denoted with a star symbol.

| **Symbol** | **Description** | **Unit** |
|---|---|---|
| *State variables and kinetic parameters* | | |
| $X_i$ | Biomass concentration | $\mathrm{g\,L^{-1}}$ |
| $P_i$ | Product concentration | $\mathrm{g\,L^{-1}}$ |
| $S_i$ | Substrate concentration | $\mathrm{g\,L^{-1}}$ |
| $\mu^{\max *}$ | Maximum growth rate | $\mathrm{h^{-1}}$ |
| $\mu_i$ | Growth rate | $\mathrm{h^{-1}}$ |
| $K_i^{\mathrm{S}*}$ | Monod constant | $\mathrm{g\,L^{-1}}$ |
| $\delta^*$ | Cell death rate | $\mathrm{h^{-1}}$ |
| $\pi_i^{0*}$ | Growth-decoupled production rate | $\mathrm{h^{-1}}$ |
| $\pi_i^{\mu *}$ | Growth-associated production rate | – |
| ${\rho_i}^*$ | Maintenance rate | $\mathrm{h^{-1}}$ |
| $Y_i^{\mathrm{X/S}*}$ | Biomass yield coefficient | $\mathrm{g\,g^{-1}}$ |
| $Y_i^{\mathrm{P/S}*}$ | Product yield coefficient | $\mathrm{g\,g^{-1}}$ |
| $Y_i^{\mathrm{ATP/S}*}$ | ATP yield coefficient | $\mathrm{g\,g^{-1}}$ |
| $\sigma$ | Substrate uptake rate | $\mathrm{h^{-1}}$ |
| *Process parameters* | | |
| $F_i$ | Feed rate | $\mathrm{L\,h^{-1}}$ |
| $V_i^*$ | Reactor volume | L |
| $D_i$ | Dilution rate | $\mathrm{h^{-1}}$ |
| $\nu$ | Volume ratio | – |
| $\phi$ | Feed split ratio | – |
| $S_i^{\mathrm{F}*}$ | Feed substrate concentration | $\mathrm{g\,L^{-1}}$ |
| $s_{2,\min}^{\mathrm{F}}$ | Minimum stage 2 feed | $\mathrm{g\,L^{-1}}$ |
| $s_{2,\max}^{F*}$ | Maximum stage 2 feed | $\mathrm{g\,L^{-1}}$ |
| *Inhibition parameters* | | |
| $X^{\max *}$ | Maximum biomass concentration | $\mathrm{g\,L^{-1}}$ |
| $P^{\max *}$ | Maximum product concentration | $\mathrm{g\,L^{-1}}$ |
| $K_i^{\mathrm{I}*}$ | Substrate inhibition constant | $\mathrm{g\,L^{-1}}$ |

*Appendix A.2. Closed-form steady-state solutions*

For the base-case model formulation, assuming Monod kinetics for substrate-limited growth, Luedeking-Piret kinetics for product formation, no substrate or product inhibition, and growth arrest in the second reactor stage, the steady-state solutions can be expressed in closed form. Setting the dynamic mass-balance equations (4) to zero yields a system of algebraic equations describing the steady state concentrations of biomass, substrate, and product. Solving these equations gives closed-form expressions for the one-stage process and two-stage process, which are used throughout the implementation described in the main text to evaluate process performance and compute $STY$:

$$X_{\mathrm{OS}} = \frac{S^{\mathrm{F}}\mu^{\max}(D - D^{\max})}{(D^{\max}+\delta)(D+\delta-\mu^{\max})\left(\frac{D+\delta}{Y_i^{\mathrm{X/S}}} + \frac{\pi^0+\pi^{\mu}(D+\delta)}{Y_i^{\mathrm{P/S}}} + \frac{m}{Yas}\right)\frac{1}{D}}, \tag{A.1a}$$

$$S_{\mathrm{OS}} = \frac{S^{\mathrm{F}}(D^{\max}+\delta-\mu^{\max})(D+\delta)}{(D^{\max}+\delta)(D+\delta-\mu^{\max})}, \tag{A.1b}$$

$$P_{\mathrm{OS}} = X_{\mathrm{OS}}\frac{\pi^0 + (D+\delta)\pi^{\mu}}{D} \tag{A.1c}$$

For the two-stage process, the steady-state solution is obtained by considering the two reactor stages sequentially. The concentrations leaving the first stage are given by:

$$X_1 = \frac{S_1^{\mathrm{F}}\mu^{\max}\phi\left(D^{\max}(\nu-1)+D\phi\right)}{(D^{\max}+\delta)\left((\delta-\mu^{\max})(\nu-1)-D\phi\right)} \tag{A.2a}$$

$$\frac{1}{\frac{\nu-1}{D}\left(\frac{\pi_1^0+\pi_1^{\mu}\delta}{Y_1^{\mathrm{P/S}}} + \frac{\delta}{Y_1^{\mathrm{X/S}}} + \frac{m_1}{Y_1^{\mathrm{ATP/S}}}\right) - \left(\frac{1}{Y_1^{\mathrm{X/S}}} + \frac{\pi_1^{\mu}}{Y_1^{\mathrm{P/S}}}\right)\phi}, \tag{A.2b}$$

$$S_1 = \frac{S_1^{\mathrm{F}}(D^{\max}+\delta-\mu^{\max})(\delta(\nu-1)-D\phi)}{(D^{\max}+\delta)((\delta-\mu^{\max})(\nu-1)-D\phi)}, \tag{A.2c}$$

$$P_1 = X_1\frac{(1-\nu)\left(\pi_1^0+\delta\pi_1^{\mu}\right)+D\phi\pi_1^{\mu}}{D\phi} \tag{A.2d}$$

The outlet of the first stage serves as the inlet condition for the second stage. The corresponding steady-state concentrations in the second reactor are:

$$X_2 = \frac{DX_1\phi}{D + \delta\nu}, \tag{A.3a}$$

$$P_2 = X_2 \frac{p_1(D + \delta\nu) + X_1 \nu \pi_2^0}{DX_1}, \tag{A.3b}$$

$$S_2 = S_2^{\mathrm{F}}(1-\phi) + \phi\nu \frac{S_1(D/\nu + \delta) - X_1\left(\frac{m_2}{Y_2^{\mathrm{ATP/S}}} + \frac{\pi_2^0}{Y_2^{\mathrm{P/S}}}\right)}{D + \delta\nu} \tag{A.3c}$$

These expressions fully define the steady-state behavior of both process configurations for any feasible combination of operating conditions and physiological parameters.

*Appendix A.3. Case studies*

*Appendix A.3.1. Model parameters and operating conditions*

Table A.2: Case-specific model parameters and operating conditions used for the PHB and lactic acid case studies.

| **Parameter** | **PHB** | **Lactic acid** |
|---|---|---|
| $\mu^{\max}$ | 0.25 | 0.48 |
| $\delta$ | $1 \times 10^{-4}$ | 0 |
| $K_i^{\mathrm{S}}$ | 1.17 | 1.2 |
| $\pi_1^0$ | 0 | 0.02 |
| $\pi_1^{\mu}$ | 0.13 | 2.20 |
| $\pi_2^0$ | 0.23 | 0.02 |
| $Y_i^{\mathrm{X/S}}$ | 0.55 | 0.40 |
| $Y_i^{\mathrm{P/S}}$ | 0.477 | 1.00 |
| $Y_i^{\mathrm{ATP/S}}$ | 1.00 | 0 |
| $\rho_i$ | 0.064 | 0 |
| $S^{\mathrm{F}}$ | 67 | 60 |
| $V$ | 9 | 1.0 |
| $s_{2,\max}^{\mathrm{F}}$ | 500 | 500 |
| $K_i^{\mathrm{I}}$ | – | 164 |
| $X^{\max}$ | – | 30 |
| $P^{\max}$ | – | 98 |

*Appendix A.3.2. PHB case study*

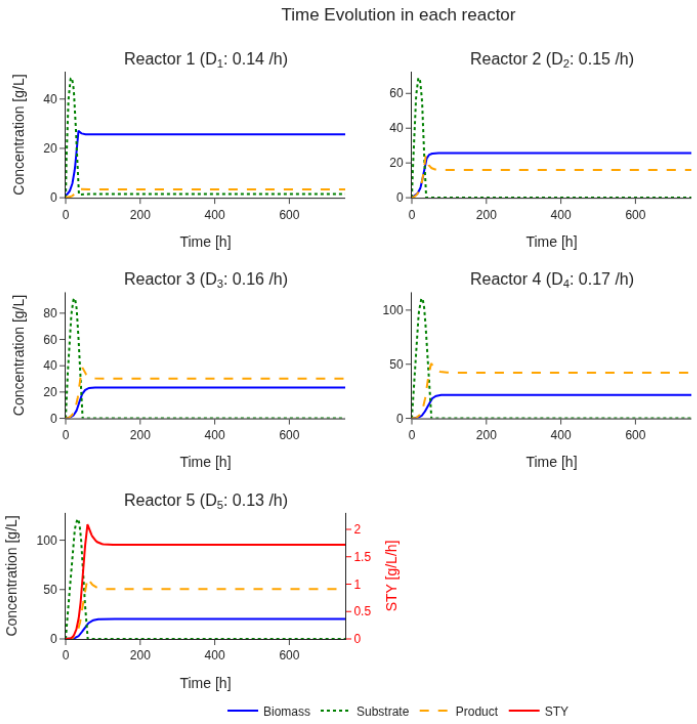


Figure A.5: Dynamic simulation of the five-stage PHB production cascade. Time evolution of biomass, substrate, and PHB concentrations in reactors R1-R5. The simulations reproduce biomass formation and downstream PHB accumulation observed in the nitrogen-limited cascade. In all reactors, substrate is depleted without accumulation, approaching zero in the downstream production stages.

| **5 reactor set-up - Experimental** | | | | | | |
|---|---|---|---|---|---|---|
| Reactor | $V_i$ L | $F_i$ $\mathrm{L\,h^{-1}}$ | $D$ $\mathrm{h^{-1}}$ | $X$ $\mathrm{g\,L^{-1}}$ | PHB $\mathrm{g\,L^{-1}}$ | $STY$ $\mathrm{g\,L^{-1}\,h^{-1}}$ |
| R1 | 1.6 | 0.222 | | 25 | 1.0 | |
| R2 | 1.6 | 0.2416 | | 27 | 15 | |
| R3 | 1.7 | 0.2639 | | 24 | 36 | |
| R4 | 1.7 | 0.2585 | | 20 | 51 | |
| R5 | 2.4 | 0.3063 | 0.034 | 20 | 63 | 2.14 |
| **5 reactor set-up - This work** ($\mu_2 = 0$, $\pi_2 = \pi(\mathrm{R3} - \mathrm{R5})$) | | | | | | |
| R1 | 1.6 | 0.222 | | 25.67 | 3.34 | |
| R2 | 1.6 | 0.2416 | | 23.57 | 18.29 | |
| R3 | 1.7 | 0.2639 | | 21.56 | 32.66 | |
| R4 | 1.7 | 0.2585 | | 20.0 | 44.82 | |
| R5 | 2.4 | 0.3063 | 0.034 | 18.55 | 53.34 | 1.81 |
| **5 reactor set-up - This work** ($\mu_2 = 0.05\mu^{\mathbf{max}}$ , $\pi_2 = \pi(R3 - R5)$) | | | | | | |
| R1 | | | | 25.66 | 3.34 | |
| R2 | | | | 25.69 | 16.02 | |
| R3 | | | | 23.51 | 30.20 | |
| R4 | | | | 21.69 | 42.22 | |
| R5 | | | | 20.23 | 50.52 | 1.72 |
| **Optimized two-stage process** | | | | | | |
| R1 | 2.43 | 0.49 | | 25.85 | 3.36 | |
| R2 | 6.57 | 0.63 | 0.07 | 19.88 | 50.28 | 3.52 |

Table A.3: Steady-state biomass (X) and PHB concentrations in the five-reactor cascade: comparison between experimental data and our model implementation. Experimental data are taken from Atlić et al. [33]. The model reproduces the observed PHB accumulation trend along the cascade, showing good agreement with experimental results under steady-state conditions.

*Appendix A.3.3. Lactic acid case study*

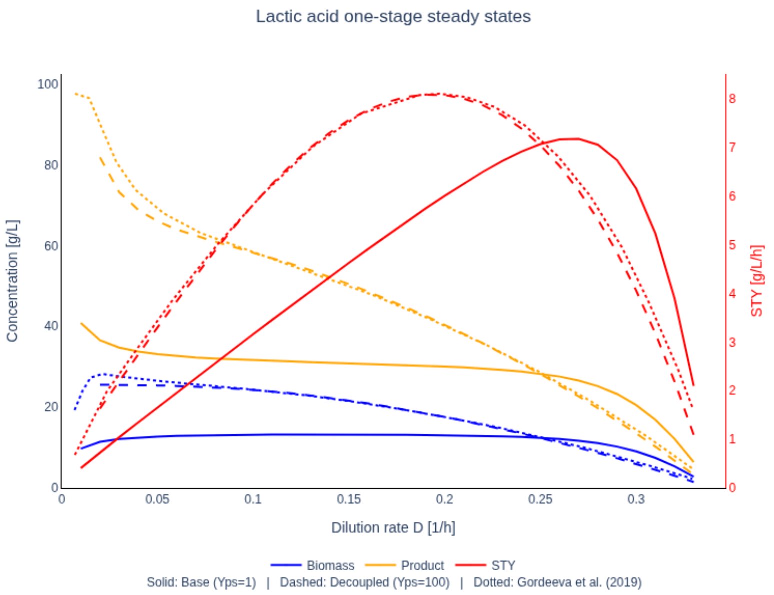


Figure A.6: Steady state profiles of biomass, product and $STY$ as a function of dilution rate for one-stage continuous lactic acid fermentation. Solid lines show the baseline configuration with $Y^{P/S} = 1\,g\,g^{-1}$, dashed lines show the decoupled case with $Y^{P/S} = 100\,g\,g^{-1}$, where substrate consumption is effectively linked only to growth. Dotted curves correspond to data from Gordeeva et al. [17]. Biomass and product concentrations are shown on the left axis, and $STY$ on the right axis. Including substrate consumption for product formation reduces $STY$ across all dilution rates and shifts the maximum towards higher dilution rates.
